# Portable low-cost macroscopic mapping system for all-optical cardiac electrophysiology

**DOI:** 10.1101/2022.04.24.489308

**Authors:** Yuli W. Heinson, Julie L. Han, Emilia Entcheva

## Abstract

**Significance:** All-optical cardiac electrophysiology enables the visualization and control of key parameters relevant to the detection of cardiac arrhythmias. Mapping such responses in human induced pluripotent stem-cell-derived cardiomyocytes (hiPSC-CMs) is of great interest for cardiotoxicity and personalized medicine applications.

**Aim:** We introduce and validate a very low-cost compact mapping system for macroscopic all-optical electrophysiology in layers of hiPSC-CMs.

**Approach:** The system uses oblique trans-illumination, low-cost cameras, light-emitting diodes and off-the-shelf components (total < $15,000) to capture voltage, calcium and mechanical waves under electrical or optical stimulation.

**Results:** Our results corroborate the equivalency of electrical and optogenetic stimulation of hiPSC-CMs, and Vm – [Ca^2+^]i similarity in conduction under pacing. Green-excitable optical sensors are combinable with blue optogenetic actuators (Chanelrhodopsin2) only under very low green light (< 0.05mW/mm^2^). Measurements in warmer culture medium yield larger spread of action potential duration and higher conduction velocities compared to Tyrode’s solution at room temperature.

**Conclusions:** As multiple optical sensors and actuators are combined, our results can help handle the “spectral congestion” and avoid parameter distortion. We illustrate the utility of the system for uncovering the action of cellular uncoupling agents and show extensibility to an epi-illumination mode for future imaging of thicker native or engineered tissues.

## 1 Introduction

Functional optical mapping of cardiac electrophysiology has become indispensable in the quantification of arrhythmia emergence and mechanisms, including in recently more widely used human induced stem-cell derived cardiomyocytes, iPSC-CMs^1–3^. Microscope-based optical mapping in cultured cells using fluorescent dyes for voltage and calcium yields acceptable signal-to-noise ratio (SNR) due to the optimized high-NA optics^4^. However, capturing wave propagation at the microscale (small field of view, FOV) is difficult as it necessitates > 5kHz temporal resolution and has been achieved exclusively using custom-built photodiode arrays^5–7^. To extend these measurements to arrhythmia-relevant spatial scales, a large FOV, closer to a centimeter, is needed to match the wavelength of the cardiac waves (action potential duration times conduction velocity). At such large FOV, optical mapping in cultured cells is much more challenging than whole-heart imaging due to difficulties in gathering enough photons from a single cell layer at high speed^4,8^. The first attempts at such large-FOV optical mapping in cultured cardiomyocytes struck a compromise between spatial and temporal resolution^9–11^. Bub et al.^11^ used lower-speed calcium wave mapping in neonatal rat cardiomyocytes to elucidate complex bursting spiral wave dynamics – a first demonstration that simple two-dimensional cardiac systems can sustain spiral waves seen at the whole heart level^12–14^. Arutunyan et al.^9^ deployed lower-speed confocal microscopy of calcium to capture ischemia-reperfusion triggered arrhythmias. We used a high-speed, lower spatial-resolution contact fluorescence imaging (CFI) of voltage with an array of photodiodes and fiber optics to directly capture the virtual electrodes induced by applied electric fields during an anatomical reentry^10^.

With the emergence of human iPSC-CMs, replacing to a large degree the neonatal rat cardiomyocyte cell culture system as an experimental *in vitro* model, interest in sensitive large-FOV optical mapping for cultured cells has increased^1,2,15^. This is dictated by the real translational opportunities to use human iPSC-CMs for preclinical studies - cardiotoxicity testing and drug discovery^16^. When combined with optogenetic stimulation, these approaches yield scalable all-optical electrophysiology solutions^17–20^. They provide powerful means for imaging and control of excitation waves^21–23^, yet, the “spectral congestion” brought about by the simultaneous use of multiple optical sensors and actuators^20,24^ requires special attention to the experimental conditions. An expensive high-end system for all-optical electrophysiology in human iPSC-CMs with large FOV was deployed by Werley et al.^25^ and has entered commercial use for drug testing. Other commercial imaging systems for high-throughput analysis, e.g., FLIPR by Molecular Devices, tend to use calcium imaging as a proxy for voltage mapping of drug responses due to the superior SNR by calcium-sensitive indicators compared to voltage-sensitive dyes. Due to challenges with sensitivity, especially in macroscopic voltage mapping in cell culture, the cameras used in optical mapping, have been traditionally high-end specialized scientific cameras, i.e., intensified CCDs, EMCCDs, and scientific CMOS (sCMOS) cameras, with a price range of $30,000 to $100,000, making them by far the most expensive part of an imaging system. Over the last decade, driven by technological improvements in mobile phone cameras, CMOS chips have undergone dramatic performance improvement, including a boost in sensitivity at a lower price point.

The goal of the present study was to build a stand-alone compact and inexpensive macroscopic (large FOV) system for all-optical electrophysiology, with capabilities for multiparametric mapping of voltage, calcium, and contraction waves in thin experimental preparations, such as layers of human iPSC-CMs. Towards this goal, we demonstrate the utility of a recent generation of machine-vision CMOS cameras, at a price that is two orders of magnitude lower than the cameras typically used in optical mapping. Along with low-cost LEDs and simple design, the total cost of the full multi-camera system is < $15,000. We perform rigorous quantification of key electrophysiological parameters across various experimental conditions, including electrical vs. optical stimulation, voltage vs. calcium mapping, various experimental solutions, and light intensities. In a companion study, we also explore in detail the links between electrical and mechanical waves tracked by such a system (Liu W et al, 2022). Overall, new insights about the impact of illumination parameters in all-optical electrophysiology are presented here that can be critical in guiding future all-optical electrophysiology studies. In example applications, we show that the system can detect conduction changes due to drugs targeting cell-cell coupling. We also demonstrate that the system, which is based on oblique transillumination in the current study, can be re-configured into epi-illumination mode for future use with thicker tissues. The presented design and results can help the more rigorous characterization of human iPSC-CMs, help streamline drug development and have overall positive impact on basic and translational studies of cardiac arrhythmias.

## 2 Methods

### 2.1 Culture of Human iPSC-Cardiomyocytes and Optogenetic Modifications

The human iPSC-CMs were the iCell2 (iCell Cardiomyocytes2 (Cat. C1016, Donor 01434) from Fujifilm/Cellular Dynamics International, Madison, WI. The hiPSC-CMs samples were cultured on fibronectin-coated (50μg/mL) 35mm dishes with 14mm glass-bottom insert (Cellvis, Mountain View, CA) at 270,000 cells per well and were grown in humidified incubator at 37 °C and 5% CO2 in manufacturer-provided culture medium and following manufacturer’s instructions. The maintenance medium was changed every other day until 7 days post-plating when measurements were conducted. For optogenetic actuation, Channelrhodopsin2 (ChR2) was expressed in the cells by adenoviral transduction with Ad-CMV-hChR2(H134R)-eYFP (Vector Biolabs, Malvern, PA) at multiplicity of infection (MOI 50) on day 5 post-plating, optimized to provide healthy and light-responsive samples^26^.

### 2.2 Functional Experiments with Human iPSC-CMs

On day 7 after plating, samples were prepared for experiments. Comparative experiments were performed either in the phenol-red containing manufacturer-provided culture medium or in Tyrode’s solution (in mM: NaCl, 135; MgCl2, 1; KCl, 5.4; CaCl2, 1.8; NaH2PO4, 0.33; glucose, 5.1; and HEPES, 5; adjusted to pH 7.4 with NaOH). Experiments in Tyrode’s solution were done at room temperature, while experiments in culture medium were performed at elevated temperature (approximately 33-35°C). Prior to optical mapping, cells were dual-labeled with spectrally-compatible fluorescent indicators for membrane voltage Vm and intracellular calcium [Ca^2+^]i in Tyrode’s solution, as described previously^18^. The [Ca^2+^]i indicator Rhod-4AM (AAT Bioquest, Sunnyvale, CA) was used at 10μM concentration for 20min, followed by a wash. Then, the near-infrared Vm dye BeRST1^27^, courtesy of Evan W. Miller (UC Berkeley), was applied at 1μM concentration for 20min. Final wash was conducted either in culture medium or Tyrode’s solution depending on experimental conditions and cells were prepared for imaging. Electrical stimulation was achieved by a bipolar platinum electrode, connected to a MyoPacer stimulator (IonOptix, Westwood, MA). Optical stimulation was integrated in the system, as described below.

In some experiments, cells were treated with cell uncoupler Heptanol (Sigma-Aldrich, St. Louis, MO) at concentration of 0.5mM for 30min. In another subset of experiments, cells were treated with an HDAC inhibitor Trichostatin A (TSA) (Sigma Aldrich)– a drug known to lead to pro-arrhythmic responses - at 100nM for 24h prior to experiments.

### 2.3 System Components

The compact macroscopic system for all-optical electrophysiology is structured on a 12” x 18” aluminum breadboard which makes it portable; all mechanical components are standard parts from Thorlabs, Newton, NJ. The system is designed for multiparametric optical imaging of Vm, [Ca^2+^]i, and dye-free (label-free) imaging of mechanical waves, under electrical and/or optical/optogenetic stimulation. Figure 1a-b shows the system, which uses two low-cost cameras, both Basler acA720-520um (Basler AG, Germany), having 720×540 pixels, 6.9μm per pixel, capable of running at up to 525 frames per second (fps). Camera 1 collects the emitted fluorescence to image Vm or [Ca^2+^]i, while simultaneously Camera 2 collects the dye-free signal visualizing the mechanical waves, see also accompanying manuscript (Liu W et al, 2022). The main imaging lens is a 50mm camera lens (0.95 F-stop, Navitar Inc. Ottawa, Canada); two additional 25mm aspherical condenser lenses (focal length 20mm, NA 0.6, Thorlabs) are placed in front of each camera to focus the image. The system is based on oblique trans-illumination for the multi-modal imaging of cardiac waves. In Fig. 1a, the bottom F1-F2-F3 cube combines the excitation light for Vm and [Ca^2+^]i imaging and directs it to the sample via oblique illumination. The top F1-F2-F3 cube represents an identical module that allows for epi-illumination regime, validated here and useful for future imaging of non-transparent samples. For oblique trans-illumination, the excitation light from the bottom cube is reflected by a mirror oriented at 30° from the horizontal plane, so that the angle of incident light related to the sample is 60°.

**Fig. 1.**
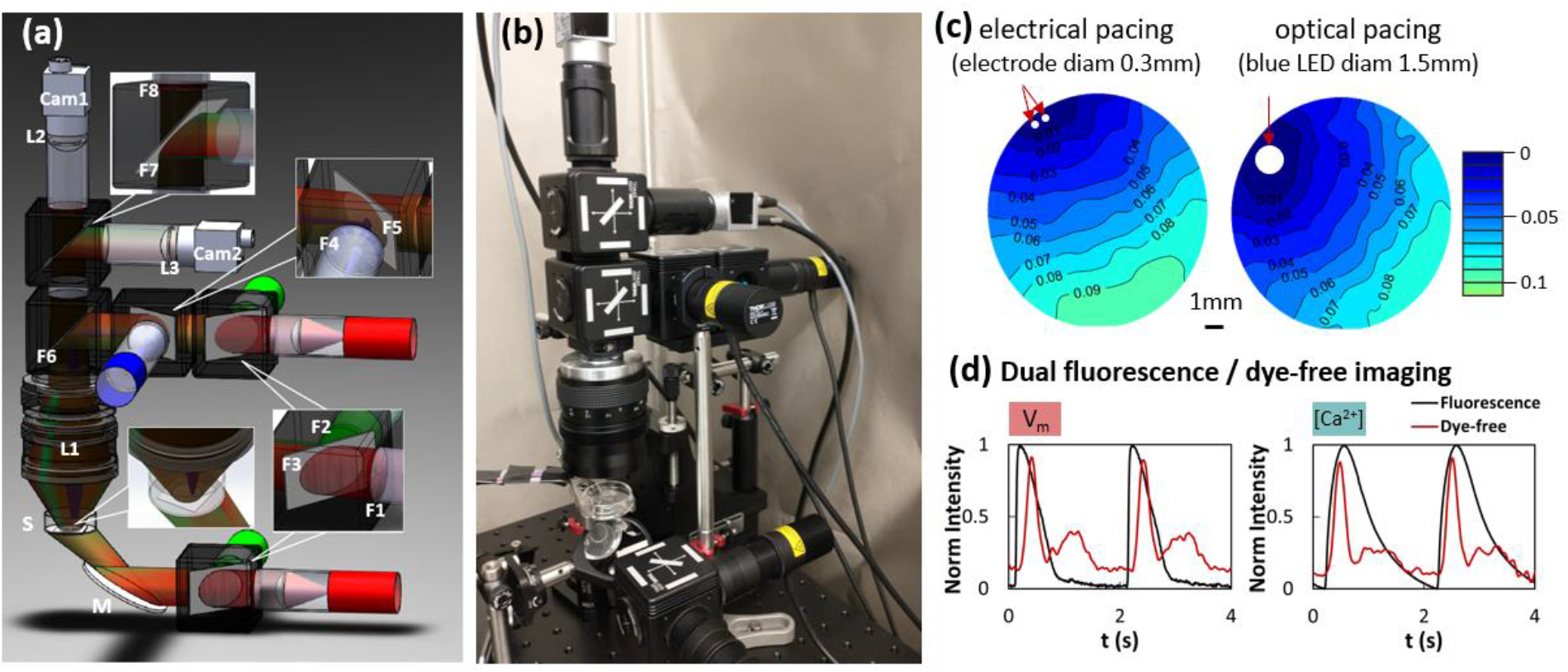
Portable low-cost all-optical macroscopic system. (a) Schematics of the imaging system generated using SolidWorks. Notations include the sample (S), CMOS cameras Cam1 (for fluorescence voltage/calcium measurements) and Cam2 for simultaneous dye-free contraction measurements. Macro-imaging lens is L1 and focusing lenses in front of the cameras are L2, L3. Blue LED is used for optogenetic stimulation, which is focused on a spot; green LED is used for excitation of calcium-responsive optical or optogenetic sensors (Rhod4, jR-GECO), and red LED for excitation of optical voltage sensors in the NIR range, e.g. BeRST1. Angled mirror (M) is used for oblique trans-illumination. Filters and dichroic mirrors include: F1: 655/40; F2: 535/30; F3: 640LP dichroic; F4: 470/24; F5: 495LP dichroic; F6: 495LP dichroic (for trans-illumination) or 473/532/660 dichroic (for epi-illumination); F7: 473/532/660 dichroic (for trans-illumination) or none (for epi-illumination); F8: 595/40+700LP. (b) Picture of the imaging system on a 12” x 18” aluminum breadboard. Total cost was < $15,000, including two cameras, three light sources and drivers, optical and mechanical components. (c) Examples of Vm activation maps at 0.5 Hz electrical pacing and optical pacing frequency, with a 1mm scale bar and a color bar indicating the activation time for the wavefront (time is in seconds). (d) Examples of fluorescence transients during dual mapping of Vm and contraction (left) and [Ca^2+^]i and contraction (right) during 0.5Hz electrical pacing. For the dye-free transients (red), the higher peak corresponds to contraction followed by a lower peak – relaxation.

The light sources in the system include a red LED (M660L4, Thorlabs) at 660nm for excitation of the voltage-sensitive dye and a green LED (M530L4, Thorlabs) at 530nm for excitation of the calcium-sensitive dye. The red and green LEDs are equipped with excitation filters: F1 (FF02-655/40-25, Semrock, Rochester, NY) and F2 (ET535/50m, Chroma, Bellows Falls, VT), respectively, and combined by a 640 long pass dichroic mirror F3 (FF640-FDi01-25×36, Semrock). Optogenetic stimulation is done from above the sample via a blue LED (M470L4, Thorlabs) at 470nm, typical pulse duration 5-10ms, controlled by TTL signals via an LED driver. The blue light is reflected by two 495 long pass dichroic mirrors (F5, F6) (FF495-Di03-25×36, Semrock) to the sample; the blue LED is slightly tilted off its optical axis and light is focused to a 1.5mm point off center of the sample.

The emission filters and dichroic mirrors in front of the cameras are as follows: F7 - a 473/532/660 multiband dichroic mirror (ZT473/532/660rpc-xt-UF1, Chroma) reflected the dye-free signal to Camera 2 and transmitted the fluorescence signal through a multi-bandpass emission filter F8: 595/40+700LP (Semrock) to Camera 1. To perform experiments with the epi-illumination module, one can simply remove the kinematic cube which holds F6 and fill the spot with the kinematic cube insert that holds F7. All dichroic mirrors are mounted in the kinematic cubes (DFM1L, Thorlabs), which makes the system modular and switching between trans-illumination module and epi-illumination module measurements - simple. Measurements of light power were done with an optical power meter (PM100D, Thorlabs).

### 2.4 Data Acquisition and Data Analysis

The camera data acquisition can be done through a free Basler acquisition software (Pylon). In this study, we used Norpix software (Norpix, Montreal, Canada) to run the cameras and guarantee synchronization. The reported experiments were done at 100 fps, full frame, and data were stored as sequence (.seq) files (Norpix). At the selected demagnification in this study, the camera-based spatial resolution of the system was 23.6μm per pixel and the FOV was approximately 17×13mm. An accompanying manuscript (Liu W et al. 2022) describes in detail the camera co-registration, the image processing and filtering steps for both the fluorescent records and the dye-free imaging. Here, we focus exclusively on the Vm and [Ca^2+^]i records obtained under various conditions. Custom software^19,28^ (Liu W et al. 2022) in Matlab, MathWorks, Natick, MA, was used to pre-process the data (baseline correction, Bartlett spatial filter, and locally weighted temporal regression filter) before event detection. Activation maps were constructed based on the detected times of activation, and the analysis software quantified the following functional parameters of interest: action potential duration at 90% repolarization, APD90, calcium transient duration at 90% recovery, CTD90, conduction velocity, CV [cm/s].

Statistical analysis to quantify the effects of various experimental conditions was done using multi-way ANOVA with Tukey post-hoc correction, and with significance considered at p < 0.05. The analysis was performed in GraphPad Prism, San Diego, CA.

## 3 Results

### 3.1 Design of a Low-Cost Portable Macroscopic All-Optical Electrophysiology System

The integrated portable low-cost all-optical macroscope is shown in Fig. 1a-b. The total cost of the system, including two cameras, all optical components, all light sources and drivers, and the mechanical scaffolding is less than <$15,000, substantially cheaper compared to existing optical mapping solutions. Key enabling technologies for building such a system are as follows: 1) improved low-cost machine-vision CMOS cameras^29,30^ over the last 5 years, offering sufficient sensitivity, suitable speed, spatial resolution and spectral response to work in the demanding low-light conditions of fast fluorescence imaging in cell culture; 2) developments in LEDs, that allow easy on-off control and provide a lower-cost, stable alternative to incandescent lamps and lasers, so that they entered the optical mapping field almost 20 years ago^31,32^; 3) improved voltage-sensitive probes in the near-infrared region developed recently, that offer sufficient SNR at high speed^27,33^; 4) development of optogenetic actuation tools, which offer optical control of cell function^20,34^.

The all-optical macroscopic system reported here includes capabilities for optogenetic actuation and multi-parametric optical mapping of Vm, [Ca^2+^]i and mechanical waves. The latter is made possible by interferometric dye-free (label-free) imaging with oblique trans-illumination, as shown by Burton et al.^21^, expanding upon a slightly different implementation reported earlier by Lee’s group^35,36^. A companion manuscript (Liu W et al, 2022) provides a rigorous look at the excitation-contraction coupling, made possible by such simultaneous electrical and mechanical mapping. The focus of this study was to demonstrate an all-optical macroscope with low-cost components and to provide detailed quantitative analysis of functional parameters from Vm and [Ca^2+^]i mapping under various conditions of optogenetic treatment, illumination power, electrical vs. optical pacing, and experimental media. Example activation maps under optical and electrical pacing, and example traces are shown in Fig. 1c.

### 3.2 Quantification of the Effects of Illumination Power on Key Electrophysiological Parameters in All-Optical Electrophysiology

Spectrally-compatible optical actuators and sensors have been combined in various studies of cardiac function over the last decade^17,18,20–22,25,37–39^. Yet, a detailed study of the effects of irradiance in the presence of multiple optical probes is missing. In Fig. 2a, we show the spectra for spectrally-compatible optogenetic actuator (ChR2) and optical sensors – Rhod-4 (calcium indicator) and Berst1 (voltage indicator). With proper selection of filters and light intensity, these have been demonstrated previously to work well together^18^. However, the ChR2 activation spectrum does extend slightly into the green region, as pointed by an arrow in Fig. 2a, so ChR2 may potentially be engaged when sufficiently strong green light is delivered.

**Fig. 2.**
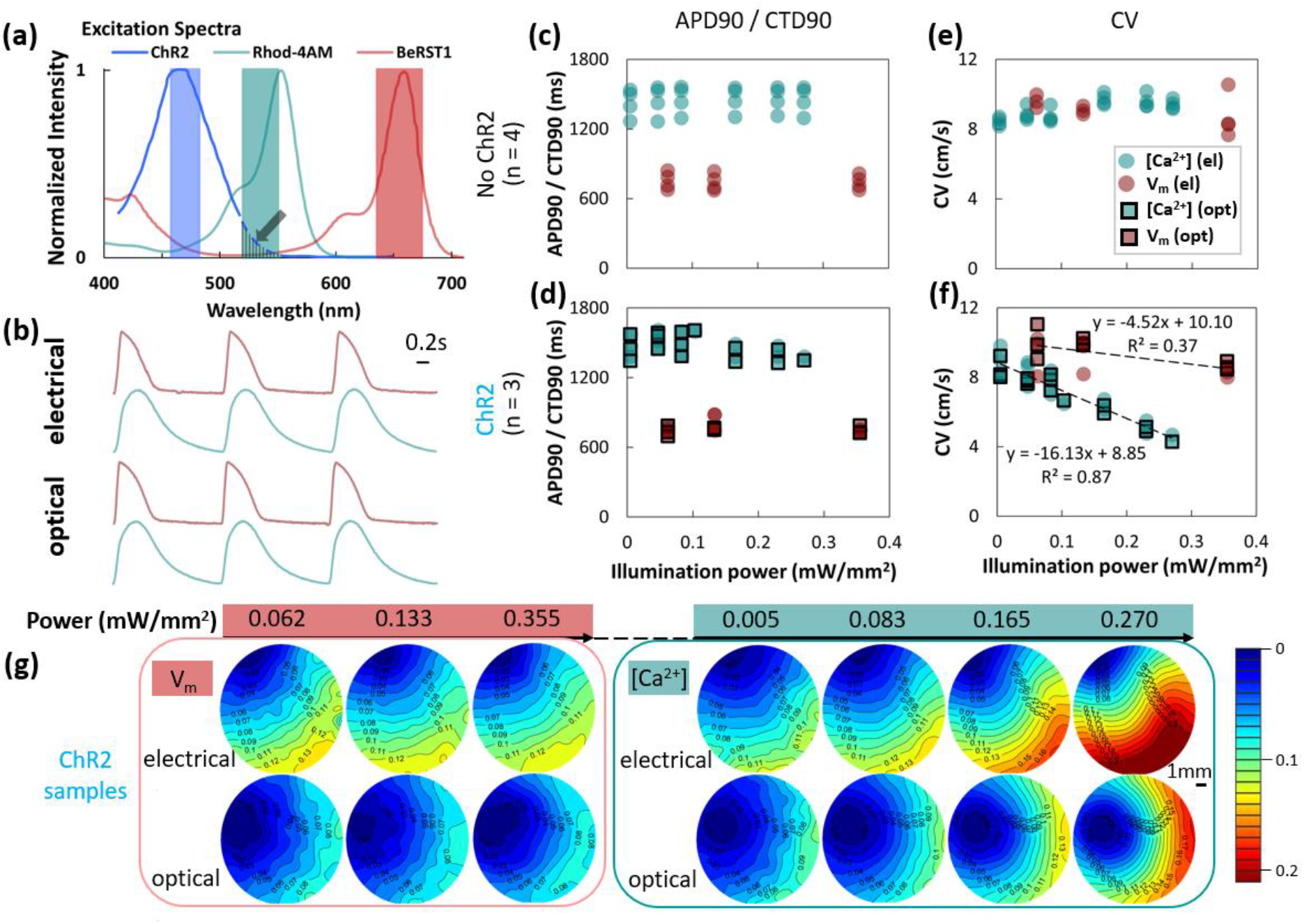
Quantifying the effect of illumination power for all-optical electrophysiology experiments in human iPSC-CMs. (a) Excitation/activation spectra of ChR2, Rhod-4AM, and BeRST1 with LED light sources filtered bandwidths shown. ChR2 spectrum has a small overlap with the green LED bandwidth (see arrow). (b) Example voltage, Vm (pink) and calcium, [Ca^2+^]i (teal) fluorescence transients 0.5Hz electrical and optical pacing. Varying illumination power (in mW/mm^2^) and keeping 0.5Hz pacing, the following parameters are shown: (c) APD90 and CTD90 of no-ChR2 samples, approximately 0.750s and 1.40s, respectively; (d) APD90 and CTD90 in response to 0.5Hz electrical or optical pacing from ChR2 samples show similar behavior to the control no-ChR2 samples. Panel (e) demonstrates constant conduction velocity, CV, from Vm or [Ca^2+^]i measurements for no-ChR2 samples for the tested illumination powers. Panel (f) shows the CVs for ChR2 samples in response to electrical and optical pacing (often completely coinciding). The CV obtained from [Ca^2+^]i mapping decreases significantly as a function of the illumination power, with a slope of -16.13, and the CV of Vm shows a very subtle decrease as illumination power increases. (g) Representative activation maps of ChR2-samples as in (f) for Vm and [Ca^2+^]i under different illumination power, with a 1mm scale bar and a color bar indicating the wavefront activation times (in seconds). Measurements were done at room temperature in Tyrode’s solution; stimulation pulses were 5ms; optical stimulation was at 1.8 mW/mm^2^. The number of samples used in each test are indicated as n. This applies for the rest of the figures.

Using the designed system, we performed a systematic study to quantify the effects of irradiance (for the excitation light in voltage and calcium imaging) on key electrophysiological parameters – APD, CTD, and CV – in all-optical electrophysiology. Example Vm and [Ca^2+^]i traces indicate equivalency of electrical and optogenetic stimulation in Fig. 2b, as suggested in earlier studies. Results in Fig. 2c-d indicate that varying red or green light power has almost no appreciable impact on APD and CTD in control or ChR2-expressing samples, as seen in prior studies^18,19^. However, the new findings here relate to the role of unintended ChR2 engagement during optical mapping and its effects on conduction velocity – Fig. 2e-f. While excitation light power did not affect CV in a significant way in control (no ChR2) samples, for optogenetically-modified samples, higher light levels significantly decreased CV in green-light obtained calcium maps only, and not in red-light obtained voltage maps (Fig. 2f-g). The effect was quite strong, such that at reasonable light levels of 0.3mW/mm^2^ for green light, conduction could be slowed by 50% when ChR2 was present. At higher light levels, engaging ChR2 by green light through the spectral cross-talk can completely block wave propagation. ANOVA analysis (Table 1) corroborates these results. Thus, safe optical mapping of calcium using green-light excitable sensors, without side-effects on conduction in the presence of an optogenetic actuator, e.g., ChR2, is only possible at very low irradiance levels, <0.05mW/mm^2^. Considering the quality of calcium sensors, these light levels still produce good SNR.

**Table 1.** P-values with two-way ANOVA analysis on illumination power and ChR2 infection for Fig. 2c-f.

| Two-way ANOVA analysis on illumination power and the presence of ChR2 |  |  |  |  |  |  |
| --- | --- | --- | --- | --- | --- | --- |
|  | APD90 / CTD90 |  |  | CV |  |  |
|  | illumination | ChR2 | illum x ChR2 | illumination | ChR2 | illum x ChR2 |
| $V_m$ | 0.6216 | 0.4493 | 0.4398 | 0.0326 | 0.6786 | 0.5770 |
| $[Ca^{2+}]_i$ | 0.6840 | 0.8846 | 0.5139 | <0.0001 | <0.0001 | <0.0001 |

### 3.3 The Impact of Experimental Conditions on Restitution and Conduction Properties for Vm and [Ca^2+^]i

Most functional optical mapping measurements in Langendorff hearts and in cardiac cell culture are done in Tyrode’s solution. For cultured human iPSC-CMs, optical measurements can be performed in culture medium especially when done within the incubator, although that is not the norm. We deployed the all-optical macroscope to quantify differences in electrophysiological parameters when the samples were studied in Tyrode’s solution (at room temperature) vs. in culture medium containing phenol-red, serum and fatty acids (at elevated temperature of about 33-35°C). Figure 3 shows the results when samples were subjected to different pacing frequencies using electrical or optical stimulation. Excitation light was kept low for calcium mapping (green LED at < 0.05mW/mm^2^) and the red LED for voltage mapping was at approximately 0.15mW/mm^2^. In all cases, in Tyrode’s and in culture medium, samples exhibited frequency adaptation (restitution), i.e., shortening in APD90, CTD90 and decrease in CV at higher pacing frequencies. CTD90 values were not influenced by the experimental conditions, while APD90 showed higher dispersion at lower pacing rates in culture medium at elevated temperature. The most dramatic effect was on conduction velocity – seen both for voltage and calcium CV, under electrical and optical pacing. Performing the experiments in culture medium at elevated temperature yielded approximately two-fold higher CVs compared to experiments in standard Tyrode’s solution at room temperature, especially at lower pacing frequencies (Fig. 3c-f). Table 2 shows the statistical ANOVA analysis of the data and indicates that in addition to the expected restitution responses, CV is significantly affected by the experimental medium and by the combination of pacing frequency and the experimental medium. Likely explanation for the increase in CV for the warmer culture medium is the strong temperature dependence of gap junctional conductance on temperature^40,41^. The use of phenol-red containing medium required adjustment (increase) in blue light for the optogenetic stimulation due to its delivery from the top of the sample, and it led to some decrease in SNR for voltage imaging, nevertheless, it was possible to obtain maps and quantification for all parameters of interest.

**Table 2.** P-values with two-way ANOVA analysis on pacing frequency and experimental conditions for Fig. 3a-d.

| Two-way ANOVA analysis on pacing frequency and experimental conditions |  |  |  |  |  |  |
| --- | --- | --- | --- | --- | --- | --- |
|  | APD90 / CTD90 |  |  | CV |  |  |
|  | frequency | solution | freq x sol | frequency | solution | freq x sol |
| $V_m$ | <0.0001 | 0.1019 | 0.0970 | <0.0001 | <0.0001 | <0.0001 |
| $[Ca^{2+}]_i$ | <0.0001 | 0.5185 | 0.9933 | <0.0001 | <0.0001 | <0.0001 |

**Fig. 3.**
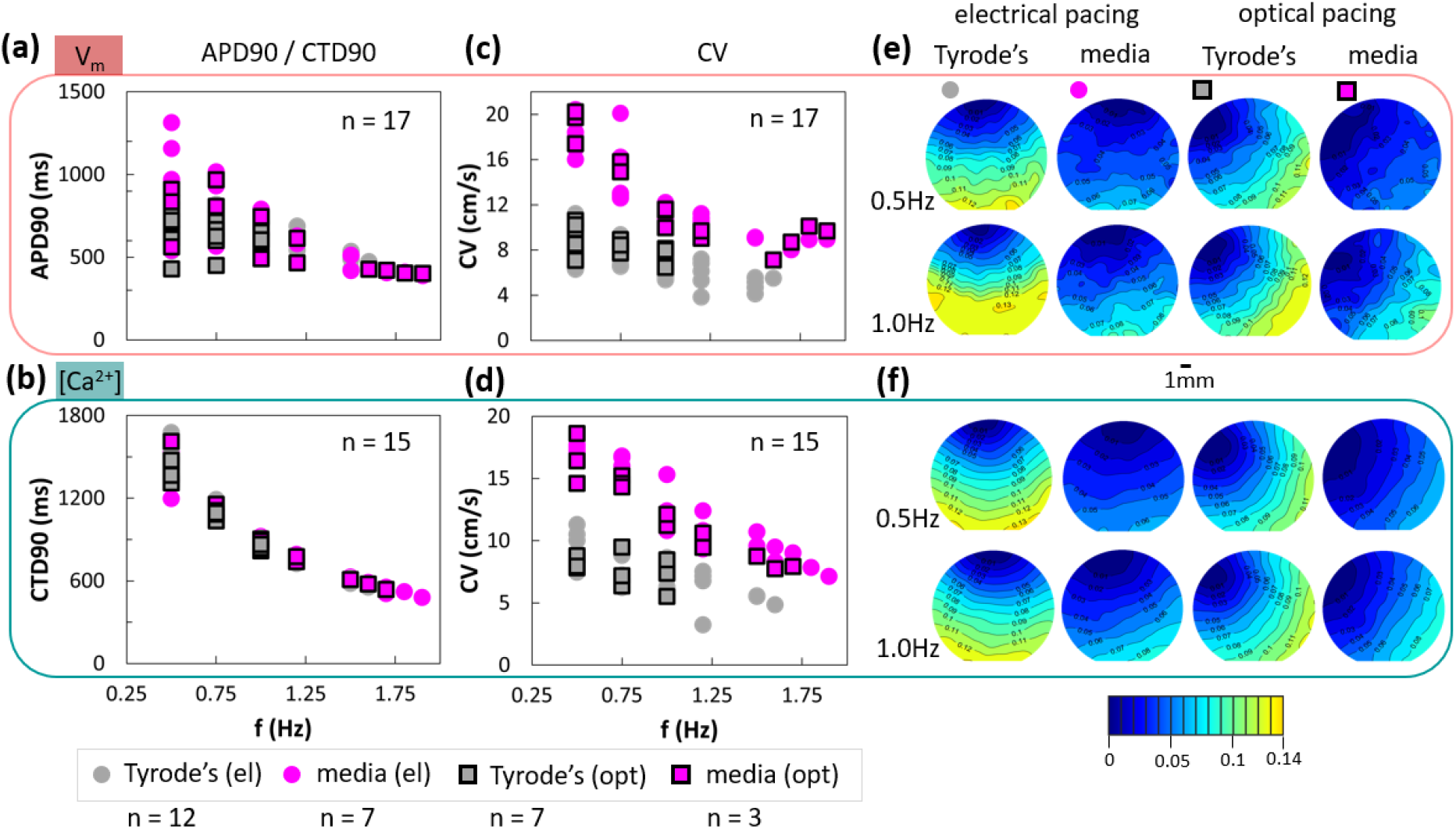
Quantifying cardiac restitution properties of human iPSC-CMs in different experimental conditions. Top - Vm results, bottom - [Ca^2+^]i results. (a-b) APD90 and CTD90 decrease with increasing pacing frequency (in Hz) as expected from adaptive restitution behavior. APD90 has a larger spread of values for lower pacing rates in culture medium at elevated temperature (33-35°C) compared to Tyrode’s solution at room temperature, while CTD90 is not very different for the two experimental conditions at any given pacing frequency. (c-d) The CVs based on Vm and [Ca^2+^]i optical mapping also show frequency-dependent decrease (restitution). Electrical and optical pacing data show similar responses. The CVs in culture medium at elevated temperature are substantially higher (almost 2x higher) compared to those in standard Tyrode’s solution at room temperature and this is true for voltage and calcium mapping under optical or electrical stimulation. Panels (e) and (f) show activation maps of representative maps as in (c) and (d), with a 1mm scale bar and a color bar showing activation times (in seconds).

### 3.4 Quantifying the Vm – [Ca^2+^]i Interrelations Under Optical vs. Electrical Pacing

While the equivalency of electrical and optical stimulation has been established when short pulses are used^42^, in Fig. 4, we present detailed comparison of APD90, CTD90 and CV in Tyrode’s and in culture medium. The correspondence of responses for all electrophysiological parameters of interest is very high (close to the identity line linking responses to electrical vs. optical stimulation). Slight divergence from the identity line is seen for high CVs, both for voltage and calcium, most likely due to the different geometry of the stimulus origin site between electrical and optical stimulation in our setup.

**Fig. 4.**
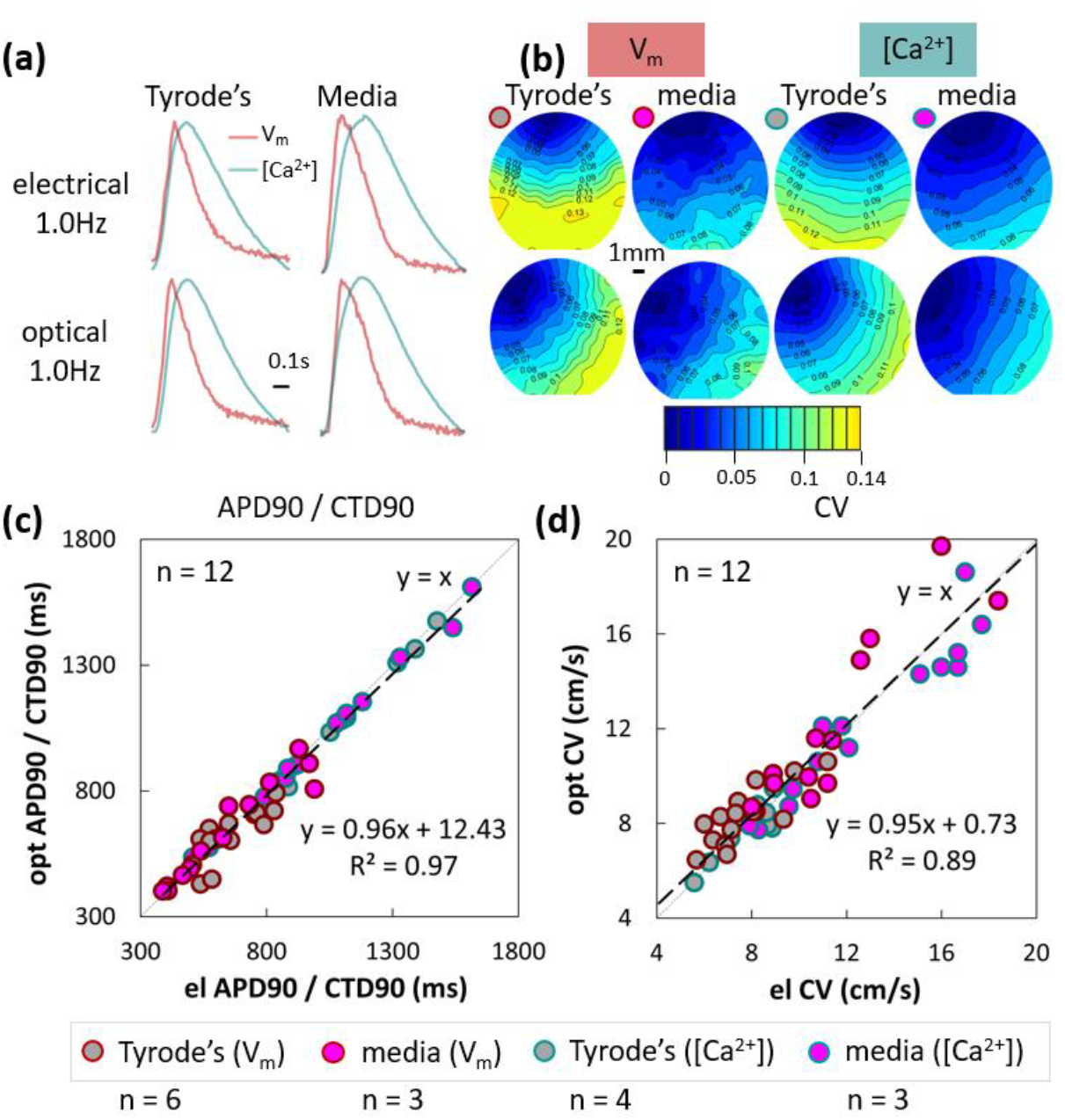
Comparison of electrophysiological responses in human iPSC-CMs to electrical vs. optogenetic pacing. (a) Fluorescence global transients for voltage and calcium at 1.0Hz electrical or optical pacing, and (b) their corresponding activation maps in culture medium or Tyrode’s solution. Panel (c) shows the APD90 and CTD90s in response to electrical and optical pacing are highly correlated (R^2^ = 0.97) and lie close to the identity line (slope of 0.96). Panel (d) shows that CVs for voltage or calcium fall on a line with a slope of 0.95 (electrical vs. optical pacing) and there is some divergence from the identity line for faster conduction in culture medium. Lightly dashed line depicts the identity line (“y=x”).

As the better SNR calcium probes are often favored in optical mapping of excitation in cell culture, it is interesting to quantify voltage-calcium relationships. Figure 5 presents APD90 vs. CTD90 in different experimental solutions and under electrical and optical pacing. As expected, APD90 is shorter than CTD90, with tendency of APD90 and CTD90 converging at high pacing rates, Fig. 5a. CVs obtained from voltage and calcium optical maps correlate well (R^2^=0.86) and fall close to the identity line (slope 1.04), regardless of the experimental medium and the mode of pacing (electrical vs. optical), Fig. 5b. Therefore, under controlled paced conditions, voltage and calcium maps provide similar information.

**Fig. 5.**
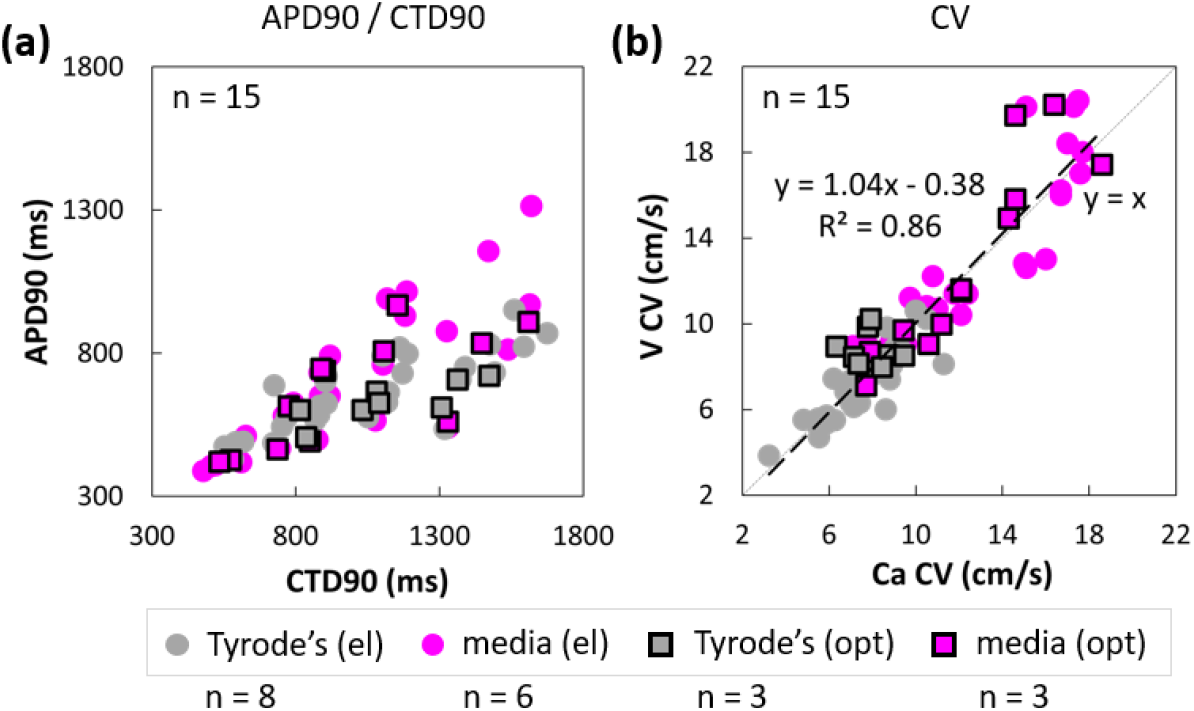
Voltage-calcium relationships in human iPSC-CMs. (a) APD90 vs. CTD90 in culture medium and in Tyrode’s solution. For shorter APD90 and CTD90 (< 0.6s, at higher pacing frequencies) the two values converge; at longer APD90 and CTD90 values, the responses are strictly under the identity line (shorter APD90 values than CTD90 values, where this relationship is exaggerated when measurements are in Tyrode’s solution. (b) CV based on voltage vs. calcium optical mapping - the fitted line has a slope of 1.04, close to the identity line (“y=x”, lightly dashed), indicating that CVs extracted from Vm or [Ca^2+^]i mapping are close to each other; some deviations are seen at high conduction velocities in culture medium.

### 3.5 Mapping the iPSC-CMs Response to Cellular Uncoupling Reagents in Different Sample Formats

The utility of the designed system was illustrated in several applications. It was applied to detect changes in cardiac electrophysiology in response to cell uncoupling drugs. In Fig. 6a-c, the responses to 0.5mM heptanol are shown (before and after drug application), under different pacing conditions, in Tyrode’s solution. As expected, at this dose, heptanol did not affect APD90 and CTD90 (Fig. 6a), however it decreased CV both obtained through voltage mapping and calcium mapping (Fig. 6b-c). ANOVA results (Table 3) confirm significant effects of both pacing frequency and heptanol on CV but not on APD90 and CTD90.

**Table 3.** P-values with two-way ANOVA analysis on pacing frequency and drug treatment for Fig. 6a-c.

| Two-way ANOVA analysis on pacing frequency and heptanol treatment |  |  |  |  |  |  |
| --- | --- | --- | --- | --- | --- | --- |
|  | APD90 / CTD90 |  |  | CV |  |  |
|  | frequency | drug | freq x drug | frequency | drug | freq x drug |
| $V_m$ | <0.0001 | 0.8492 | 0.8888 | 0.0182 | 0.0028 | 0.9956 |
| $[Ca^{2+}]_i$ | <0.0001 | 0.7599 | 0.5920 | 0.0001 | <0.0001 | 0.6949 |

**Fig. 6.**
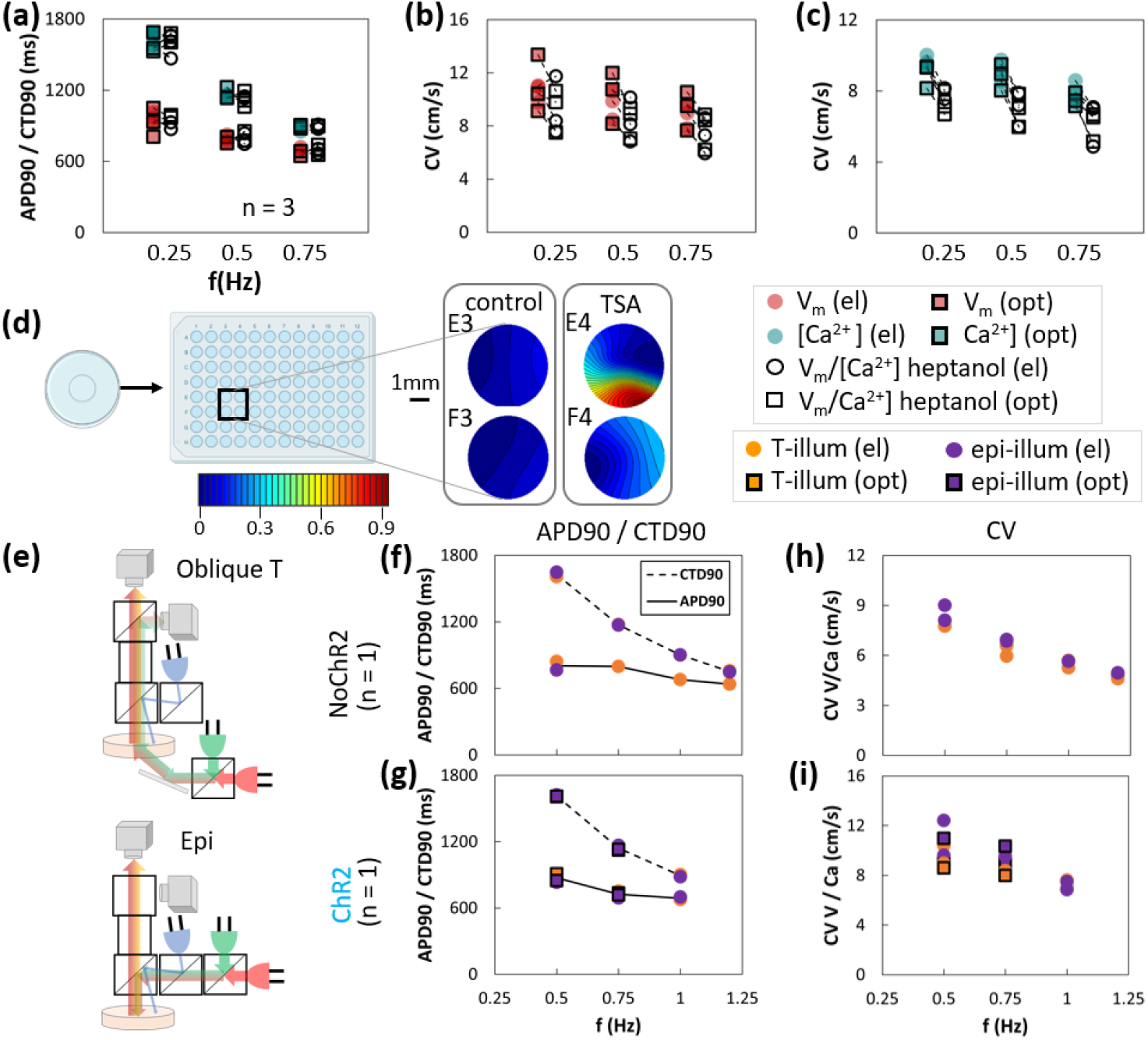
Applications and extensions of the all-optical mapping system. (a - c) Mapping changes in electrophysiological parameters (voltage and calcium) in response to treatment with a cellular uncoupling agent (30min of 0.5mM heptanol) under different pacing frequencies. (a) Heptanol had no effect on APD90 and CTD90 regardless of pacing frequency under optical or electrical stimulation. (b - c) CVs for both Vm and [Ca^2+^]i were significantly reduced upon heptanol treatment at each pacing frequency, for both electrical pacing and optical pacing (p < 0.0001 for heptanol effects based on ANOVA). (d) Measurements of responses to treatment with Trichostatin A (TSA), 100nM for 24h, in a 96-well plate, where 4 wells are imaged simultaneously (E3 and F3 - control samples and E4 and F4 - TSA-treated). Calcium activation maps are shown, where TSA appears to reduce CV; TSA induced spiral waves in a very small spatial scale in E4. (e - i) Validation of the extension of the mapping system to an epi-illumination-based imaging. (e) Schematic illustrations of the oblique trans-illumination vs. epi-illumination mode. (f - g) APD90 and CTD90 results under different pacing frequencies for control no-ChR2 samples and ChR2 samples, respectively. Orange represents trans-illumination and purple – epi-illumination. Note that there is overlap of data points from the two regimes. (h - i) CV results for voltage and calcium under different pacing conditions for no-ChR2 and for ChR2 samples, respectively. For each studied condition, restitution properties are visible. The results from trans-illumination module and epi-illumination module almost completely coincide, indicating the epi-illumination module was functional and can be deployed to measure thicker non-transparent tissue samples.

Although most experiments were done in glass-bottom 35mm dishes, in Fig. 6d, we demonstrate that in the current configuration and for the FOV approximately 19×15mm (26.9μm per pixel), the system is compatible with other formats and can map 4 wells (from a 96-well plate) simultaneously. A different drug with effects on coupling and overall electrophysiology – an HDAC inhibitor, Trichostatin-A (TSA) was applied for 24h at 100nM concentration. Conduction slowing was documented in the TSA-treated samples, as seen in the activation maps. In one of the shown samples (E4), the conduction was slowed down sufficiently to sustain a rotating wave within a very small space (each well is 7mm in diameter).

### 3.6 Validation of Extensibility to Epi-illumination

The oblique trans-illumination mode of imaging in the presented system works well with cultured iPSC-CMs, i.e., transparent monolayer samples. For the study of thicker, non-transparent samples in the future, we explored the possibility to reconfigure the system to an epi-fluorescent mode. Due to the modular design, this reconfiguring was straight-forward. Fig. 6e illustrates the difference between the oblique trans-illumination and the epi-illumination configuration. We tested out how these two systems compare in terms of measured electrophysiological parameters. We quantified APD90, CTD90, and CV for both no ChR2 and ChR2 expressing samples at various pacing frequencies, using both modules, and compared the results. Control (no ChR2) results for APD90, CTD90, and CV are shown in Fig. 6f, h; and results for ChR2-expressing samples are shown in Fig. 6g, i. All measurements in Fig. 6f–i show the characteristic restitution for both electrical and optical pacing. Note that in Fig. 6h and 6i the CVs of Vm and [Ca^2+^]i are not distinguishable from each other due to them having similar values. Without further optimization, we found that the epi-illumination mode tended to produce noisier Vm signals, which led to the corresponding APD90s to have larger uncertainties, especially at higher pacing frequencies. Overall, the epi-illumination results agree well with the trans-illumination, which verified that the epi-illumination configuration was functional and can be deployed in the future to study non-transparent samples, such as heart slices or whole hearts.

## 4 Discussion

Efforts have been put in place to standardize cardiac electrophysiology experiments by enforcing certain rules of reporting^43^. Detailed experimental descriptions and adherence to standard experimental approaches are viewed as a way towards easier comparison of findings obtained by different laboratories. Yet, the complexity of experiments in cardiac electrophysiology, as an evolving field, using a variety of experimental models, variety of configurations and reagents, and with the adoption of new technologies, e.g., novel optical tools^3^, often prevents comprehensive analysis and consideration of all aspects that may impact the results.

Here, we report on the design and validation of a portable low-cost all-optical macroscopic mapping system (Fig. 1) to study cardiac electrophysiology in human iPSC-CMs. We show that the system can also be easily reconfigured to epi-illumination mode to perform all-optical mapping in non-transparent thicker samples, such as cardiac tissue slices or whole hearts, Fig. 6e-i. The hope is that simpler and more affordable mapping systems, offering a comprehensive, multiparametric view on cardiac responses to various stimuli and perturbations, can help quantify and standardize these complex measurements. Optical mapping offers a non-invasive way to probe electrophysiological function in a multicellular context, without cell isolation, which can adversely impact function.

While optical mapping has been a valuable tool in the cardiac field for over 30 years^44^, and in the last 10 years it has been augmented with optical/optogenetic stimulation capabilities^20^, very few studies have dealt with baseline characterization of the effects of experimental conditions. For example, Kanaporis and colleagues^45^ provided rigorous analysis of the effects of illumination power in optical mapping on key assessed parameters, e.g., APD and CV. They found that APD can be shortened and CV can be increased with higher excitation light irradiances. The effects were wavelength-dependent and likely related to transient thermal responses than phototoxicity of the sensing probes. The current study uses irradiances substantially lower (in all cases < 0.5mW/mm^2^) than the ones that led to thermally-mediated change in electrophysiological parameters, as seen in^45^. Klimas et al.^18^ showed that irradiance of the excitation light can have non-linear, non-monotonic effect on the SNR in voltage and calcium imaging. Yet, very few studies quantify, report, or seriously consider these basic conditions of optical mapping experiments. When optogenetic stimulation is integrated with optical mapping using various sensors (for all-optical electrophysiology), and when their spectral profiles are close together, there is even more pressing need to carefully consider the conditions of such experiments. The current study specifically draws attention to potential unintended engagement of ChR2 actuation through the excitation light for optical sensing, when it is in the green part of the spectrum. We show that green-excitable Rhod-4 calcium sensing in optogenetically-modified cardiomyocytes only works without distortion of the results when the irradiance is very low (< 0.05mW/mm^2^), Fig. 2. Engagement/opening of ChR2 ion channels directly contributes to depolarization of the membrane and inactivation of the sodium channels, hence impacting conduction. The effect is likely not specific to Rhod-4, but extends to other optical dyes (FluoVolt, di4-ANNEPS) and genetically encoded sensors, e.g., jR-GECO^46^, excitable in the same green wavelength range. In addition to reducing the excitation light as a solution to this problem, using blue-shifted opsins, such as CheRiff^47^ may help. In general, the all-optical mapping system developed here can help characterize the concurrent deployment of a large range of optical and optogenetic sensors and actuators, as seen in use with human iPSC-CMs^48–51^.

Adding optical stimulation in all-optical mapping experiments is appealing because of its general equivalency to electrical stimulation^42^ and the many benefits it brings – contactless, spatially-resolved, cell-specific actuation^22^. Our results here corroborate the safe use of optogenetic stimulation (with short pulses) as an alternative to electrical stimulation, Figs. 3 and 4, with full preservation of APD, CTD, CV, and restitution responses. These findings are particularly impactful when considering high-throughput format testing with human iPSC-CMs for cardiotoxicity or drug development^17,18^. In such format, it is much easier to implement contactless spatially-distributed optical stimulation (rather than distributed electrical stimulation in each well) to expand the testing from observations of spontaneous activity to frequency-dependent responses when pharmacological or other therapies are tested. The system can be used in culture medium at elevated temperature (Figs 3-5) for potential long-term recordings of activity in human iPSC-CMs. As suggested in Fig. 6d, high-throughput format plates can be accommodated in this system, with appropriate further expansion of the FOV. Such scaling up in FOV has been done in industrial-grade expensive all-optical systems^25^ (although the FOV there was almost 10x smaller than the reported here), or in development of an optical mapping system of spontaneous activity within multi-well plates^52^; an alternative interesting solution for rapid mapping of multiple locations/samples has been proposed as random access dye-free imaging^53^. Examples of the use of the low-cost machine-vision cameras in complex imaging solutions at the whole heart level – e.g., for panoramic^29^ or volumetric assessment^30^ of electrophysiology – promise emergence of more affordable tools across experimental models and scales. In general, portable low-cost imaging systems are of interest in space-constrained and/or low-resource conditions, such as laboratories in the developing countries, or in mobile deployments (on the field, in space etc).

These developments, when combined with human iPSC-CMs have high translational value for pre-clinical testing of new drugs and for personalized medicine. Multi-parametric characterization of function can better inform drug risk during the preclinical evaluation. In the current study, voltage and calcium mapping was done sequentially in the same dual-labeled samples, with the possibility for simultaneous mechanical wave mapping (Liu W et al., 2022). In future studies, we intend to combine the two measurements using temporal multiplexing onto the same camera, as we have done in a microscopic version of all-optical electrophysiology^18^. Comprehensive electro-mechanical responses obtained in human iPSC-CMs over space and time under various pacing conditions can be used to develop better in silico tools for prediction of drug action^54,55^.

## Acknowledgements

The authors thank Dr. Wei Liu for developing the current version of the analysis software and PhD student Weizhen Li for help with some experiments, notably the TSA treatment of cells. This work was supported in part by grants from the National Science Foundation (PFI 1827535 and EFMA 1830941) and the National Institutes of Health (R01HL144157).

## Code, Data, and Materials Availability

The datasets from this study are available upon request to the corresponding author. All data points are explicitly presented in the Figures.

## Yuli W. Heinson

is a postdoctoral associate at George Washington University. She received her BS, MS, and PhD degrees in physics from Harbin Normal University, China in 2010, Creighton University, Omaha, NE in 2012, and Kansas State University, Manhattan, KS in 2016, respectively. Her research interests lie in the design and development of optical instruments.

## Julie L. Han

is a PhD student at George Washington University. She holds a BS degree in Chemistry from Bryn Mawr College. Her prior research was on green chemistry methods and signaling pathways in cancer. Her doctoral dissertation focuses on new optical, optogenetic and gene modulation (CRISPRi/a) methods for control of cardiomyocytes.

## Emilia Entcheva

is a Professor of Biomedical Engineering and directs the Cardiac Optogenetics and Optical Imaging Laboratory at George Washington University. Professor Entcheva is a lifetime SPIE and Optica Member and an AIMBE Fellow. Her group has been involved in the extension of optogenetic methods to cardiac applications, their integration with optical mapping techniques, and the deployment of all-optical electrophysiology in human stem-cell derived cardiomyocytes.

